# MAAPE: A Modular Approach to Evolutionary Analysis of Protein Embeddings

**DOI:** 10.1101/2024.11.27.625620

**Authors:** Xiaoyu Wang, Heqian Zhang, Jiaquan Huang, Zhiwei Qin

## Abstract

We present MAPPE, a novel algorithm integrating a k-nearest neighbor (KNN) similarity network with co-occurrence matrix analysis to extract evolutionary insights from protein language model (PLM) embeddings. The KNN network captures diverse evolutionary relationships and events, while the co-occurrence matrix identifies directional evolutionary paths and potential signals of gene transfer. MAPPE overcomes the limitations of traditional sequence alignment methods in detecting structural homology and functional associations in low-similarity protein sequences. By employing sliding windows of varying sizes, it analyzes embeddings to uncover both local and global evolutionary signals encoded by PLMs. We have benchmarked MAAPE approach on two well-characterized protein family datasets: the Als regulatory system (AlsS/AlsR) and the Rad DNA repair protein families. In both cases, MAAPE successfully reconstructed evolutionary networks that align with established phylogenetic relationships. This approach offers a deeper understanding of evolutionary relationships and holds significant potential for applications in protein evolution research, functional prediction, and the rational design of novel proteins.

## Introduction

Artificial intelligence has achieved breakthroughs in the field of protein science. The success of AlphaFold2 (1) marked a significant advancement in protein structure prediction, in the meantime large-scale language models like ESM-2 (2)and ProtGPT2 (3)have established novel paradigm for understanding of the complex interplay between protein sequences, structures, and functions. They not only predict protein structures with high accuracy but also possess profound capabilities in understanding protein evolution, function, and interactions. This milestone opens new opportunities in protein drug development, mutation effect prediction, and industrial enzyme design, innovated in life sciences and offering new solutions to major challenges in human health and environmental protection.

The Evolutionary Scale Modeling (ESM) series is among the leading models in the current landscape of protein language models. It employs a Transformer-architectured and self-supervised learning to dissect relationships between amino acid residues from billions of natural protein sequences. The latest iteration, ESM-3, is a multimodal generative language model containing up to 98 billion parameters and trained on a dataset encompassing 2.78 billion natural proteins (4). This model encodes three-dimensional structural information using discrete tokens and incorporates invariant geometric attention mechanisms, achieving comprehensive feature extraction by effectively representing protein as vector embeddings and enabling the generation of proteins.

Recent studies have revealed that protein language models (PLMs) can capture complicated evolutionary information (5). It demonstrates that language models can predict the evolutionary dynamics of proteins, the embedding space of these models reflects evolutionary distances within protein families, and even reconstructs evolutionary histories. Specifically, ESM-3 successfully generated a novel green fluorescent protein (esmGFP) with 58% sequence divergence from existing fluorescent proteins, a degree of difference comparable to that accumulated over 500 million years of natural evolution (4). This achievement demonstrates the ability of PLMs not only to comprehend the evolutionary information inherent in sequences but also to simulate and reproduce complex evolutionary processes.

Traditional sequence alignment methods have long troubled with the “twilight zone” of protein sequence similarity, where sequence identity falls below 20-35% (6). Methods such as BLOSUM matrices struggle to capture evolutionary relationships in these low-similarity regions because proteins may retain similar three-dimensional structures and functions despite significant sequence divergence (7, 8). Studies indicate that proteins with as low as 20% sequence identity can still exhibit homology and structural similarity, yet these critical evolutionary insights are often lost in sequence comparison analyses (9).

To address the difficulties of aligning low-similarity sequence regions and extracting evolutionary information, here we present the <u>M</u>odular <u>A</u>ssembly <u>A</u>nalysis of <u>P</u>rotein <u>E</u>mbeddings (MAAPE) algorithm, designed to extract evolutionary insights from protein language model embeddings. MAAPE comprises two core components: (1) a K-nearest neighbors (KNN) similarity network based on Euclidean distance, which captures various evolutionary relationships and events, including functional and structural changes, point mutations, recombination events, gene duplication, and horizontal gene transfer (HGT), etc.; and (2) a co-occurrence matrix analysis system that compares the similarity and assembly directions of subvectors across different window sizes, revealing the directional paths of evolution and signals of gene transfer. MAAPE innovatively integrates a Euclidean distance based KNN similarity network with multi-scale co-occurrence matrix analysis, enabling the capture of traditional sequence similarities while also indicating evolutionary directions. By employing sliding windows of varying sizes to analyze embeddings, MAAPE can detect local and global evolutionary signals captured by PLMs. This algorithm not only facilitates a deeper understanding of evolutionary relationships among low-similarity protein sequences but also uncovers functional associations that conventional methods might overlook, and therefore holds substantial promise for applications in protein evolution research, functional prediction, and the design of novel proteins.

## Results

We are introducing here a new algorithm named MAAPE to address the limitations of sequence alignment in structural homology and to decipher the semantic information encoded in PLM embeddings (Figure 1). MAAPE combines a k-nearest neighbour KNN similarity network that is based on the Euclidean distances between the embedded vectors and a co-occurrence matrix that measures both the similarity between the embeddings and their assembly directions (10, 11). The undirected KNN graph encapsulates various evolutionary relationships and events, such as functional and structural changes, point mutations, recombination events, gene duplications, and HGT events (12), but lacks information about the directionality of the evolutionary paths and the detection of gene transfer signals. For this purpose, we dissected embeddings with windows of different sizes, compared the Euclidean distances between subvectors of the same size, and evaluated their similarity by constructing a co-occurrence matrix. Additionally, we identified the containment relationship paths between subvectors of different sizes, assigning directionality to the matrix based on how smaller subvectors assemble into larger ones.

**Figure 1.**
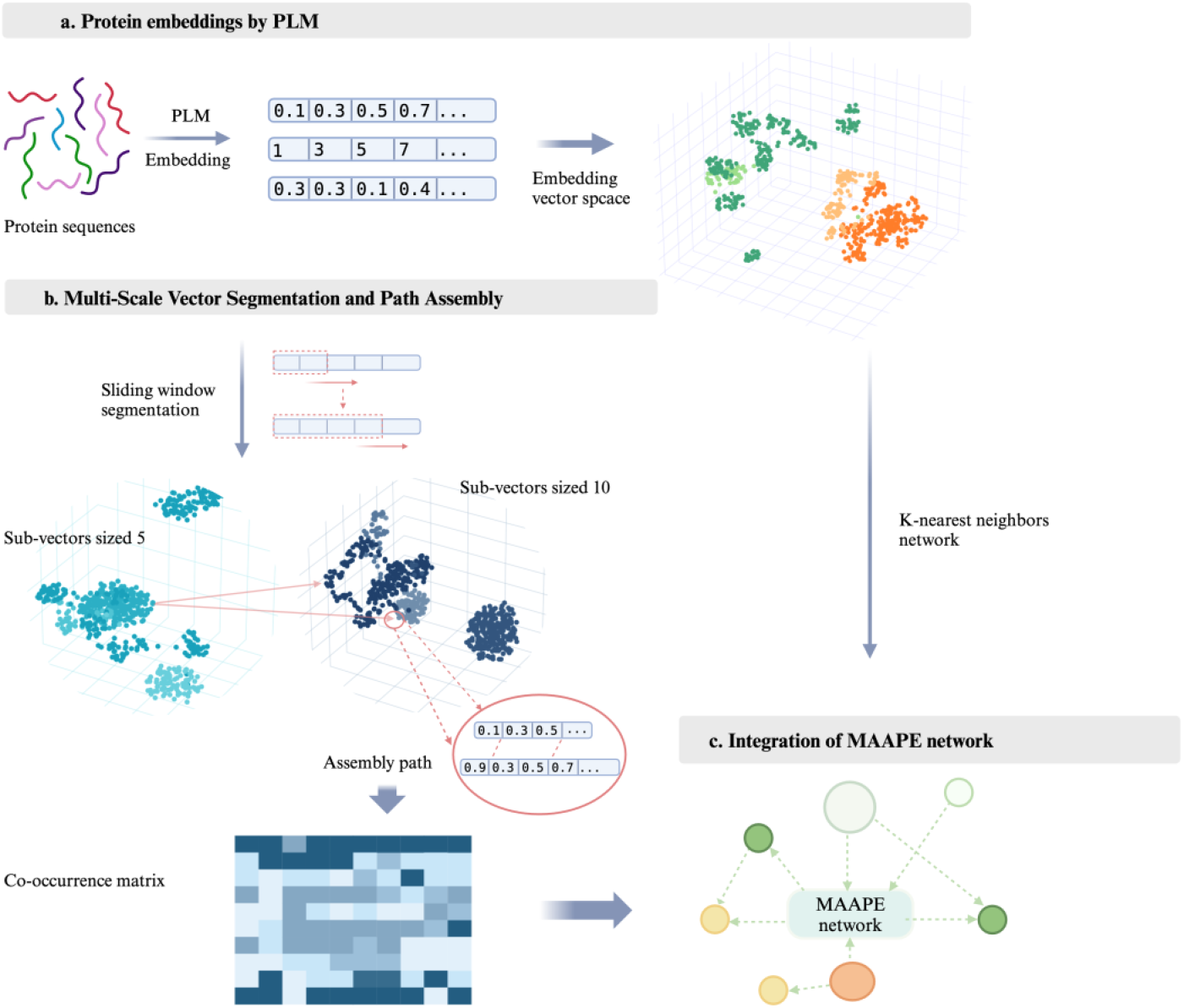
MAAPE network construction pipeline. (a) A pretrained language model (the pretrained ESM-2 model in this work) was used to transform protein sequences into high-dimensional vector representations. A UMAP-based dimensionally-reduced visualization of the embedding vector space is illustrated. (b) Multiscale vector segmentation and path assembly were implemented by the following steps. Sliding windows were employed to segment the embedding vectors, generating subvectors of different sizes (a set of small to large windows, e.g., with sizes of 5 and 10); the subvector assembly paths were connected by using Euclidean distance as the metric, where the path directions were designated from small-window vectors to large ones, and an example of an assembly path is demonstrated; a co-occurrence matrix was generated between the original vectors connected by paths, reflecting the association strengths between the vectors; and a K-NN network reflecting similarity was constructed. (c) Co-occurrence matrix information was incorporated into the KNN network to generate the MAAPE network. The relationships between different protein sequences are shown.

### Language model and feature extraction

ESM-2, based on the Transformer architecture, is specifically optimized for protein sequences and captures complex sequence patterns through self-attention mechanisms. The model was trained using a masked language modelling approach, where certain amino acids in the sequence are randomly masked. The model is trained to predict these masked positions, thereby learning rich contextual representations of the sequences. In this study, we utilized Facebook’s pre-trained ESM2_t36_3B_UR50D model to embed protein sequences, which comprise 36 layers, each with a hidden state dimension of 2560, totaling approximately 3 billion parameters. By reading protein sequences from both target and outgroup sequences, the ESM-2 tokenizer was utilized to encode these sequences, and the encoded sequences were processed in batches through the ESM-2 model. We extracted the output from the last hidden layer as feature representations of sequences (2).

For each batch *S* containing *n* sequences, where *S* is defined as *S* = {*S*_1,_ *S*_2,_ …, *S*_*n*_} and the hidden state dimension is 2560, the hidden state matrix *H* of the final layer can be expressed as *H* ∈ ℝ^*n*×*L*×2560^, where *L* is the length of the sequences. For the *i* -th sequence *S*_*i*_, we extract *E*(*S*_*i*_) = *H*_*i*_ [:, 0, :] as the embedding representation.

To further optimize the representation of the embedding vectors, we normalize all vectors to the unit hypersphere by L2 normalization on the embedding vectors. This eliminates the influence of vector length and makes similarity calculations more accurate.

### Feature dimensionality reduction

Considering the high dimensionality of the embedding vectors conduces demanding need in computing power and storage, we employ Uniform Manifold Approximation and Projection (UMAP) to reduce the dimensionality of the embedding vectors. UMAP is a nonlinear dimensionality reduction technique that is well-suited for preserving the global and local structure in high-dimensional data (13). By analysing the low-dimensional embedding through UMAP, we decided to reduce the dimension of each embedded sequence to 200, which balances computational efficiency and representation fidelity by preserving more than 99.99% of information. Therefore, we performed dimension reduction on the normalized embedding vectors to obtain a low-dimensional representation that captures the essential features of the original high-dimensional data.

### Modular transfer matrix computation of sequence embeddings

#### Sliding window slice of embedding vectors

When normalized vectors are segmented using sliding windows of varying sizes, we can identify recurrent fragments within windows of the same size. According to assembly theory (14), these recurrent fragments from the same window sizes represent sequence modules that recur across different sequences, suggesting they possess significant evolutionary/function/structural value. However, the hierarchical nesting of fragments from larger to smaller window sizes, on the other hand, can infer evolutionary trajectories where smaller modules assemble into larger ones. Therefore, the choice of window sizes is crucial. To avoid the inclusion of irrelevant information that could introduce excessive noise and obscure the signals of genuine modules, we employed the concept of information entropy to determine which window sizes provide the most informative content on average.

#### Information Entropy

Information entropy, introduced by Claude Shannon, quantifies the uncertainty or unpredictability in a set of data (15). In the context of our study, entropy measures the distribution of sequence fragments within each window. Higher entropy values indicate a more uniform distribution, suggesting a higher degree of variability and, by extension, more valuable information. Conversely, lower entropy values suggest redundancy and less informative content. By calculating the average entropy across different window sizes, we can objectively identify which sizes yield the most informative and potentially significant fragments, and entropy was computed using the formula:

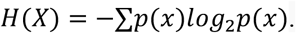

Upon this approach, we selected window sizes that maximize average information entropy, stride is set to 1 to preserve context information learned by the language model, thereby enhancing our ability to detect meaningful evolutionary modules. The chosen window sizes are as follows: [5, 10, 15, 20, 25, 30, 35, 40, 45, 50, 60, 70, 80, 90, 100, 120, 140, 150, 200].

#### Pairwise similarity search for vector fragments

We utilized the IndexFlatL2 index of the Faiss library to perform a similarity search on the split vector fragments for each window size, those with Euclidean distances less than a threshold were set to be recurrent modules and saved for downstream analysis (16). To determine a proper threshold for each window, we calculated the pairwise Euclidean distances for each window size and plotted a histogram to visualize the distribution of these distances, identified the bin whose range was closest to 0, the lower bound of this closest bin was selected as its threshold, in vectors with other window sizes, use the square root transformation based on the corresponding vector length. This bin represents the smallest distances in the distribution of the smallest vector slices, this excludes other influencing factors, only indicative of highly similar vector fragments.

#### Modularly transfer matrix computation

We further explored the modular transfer routes between sequences. Starting from the smallest fragments, we searched for their containment relationships within the next-level fragments by calculating their Euclidean distance and iterated this process to the largest vectors. Within each search cycle, we define the containment relationship found as the process of passing assembled modules between sequences, i.e., how complex sequential modules evolve from simpler ones through horizontal transfer, etc., and we define the direction of subordination from small to large vectors as the direction of assembly trajectories.

An intuitive consideration is that smaller modules are easier to form compared to larger ones, thus can be observed more often. Therefore, in our subsequent calculations, we assign larger weight coefficients to larger vectors according to their occurrence frequency.

A co-occurrence matrix between the original sequences is calculated from containment relationships under different window-sized vectors, which reflects the proximity and transfer direction between target sequences. For target sequences *S*1, *S*2, …, *Sn*, where *n* is the number of sequences, for each window size *w*, the co-occurrence matrix is defined as:

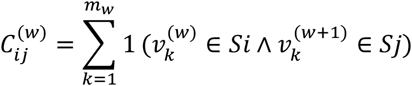

Where the element 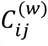 represents the co-occurrence count of the containment vectors between vector 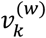 from sequence *Si* and 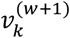 from sequence *Sj, mw* is the number of containment vectors for window size *w*, and 1(·) is the indicator function.

For each window *w*, a weight matrix *W*^(*w*)^ is defined as element 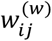 represents the edge weight between sequences *S*_*i*_ and *S*_*j*_ for the window size *w*:

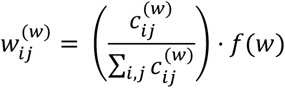

Where 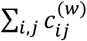 is the total co-occurrence count for the window size *w*. The function *f*(*w*) is a function that increases with window size to assign higher weights to larger vectors.

The directions for each pair of sequences *S*_*i*_ and *S*_*j*_, is calculated by comparing the forward edge weight 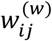 and the reverse edge weight 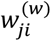:

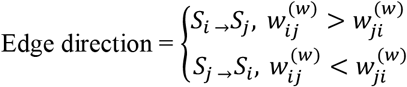

The weight of the edge between *S*_*i*_ and *S*_*j*_ is the sum of the weights in both directions:

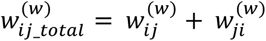

A directed acyclic graph was constructed through the co-occurrence weight matrix, and edges within the upper 20% quantile were kept for visualization.

The edge weights in the graph contain information about both the sequence similarity represented by the co-occurrence counts and the assembly direction from small vectors to large vectors, providing valuable insights for understanding the evolutionary patterns of protein sequences.

### KNN graph for sequence similarity representation and evolution trajectory integration

#### KNN graph construction

Although our task of modular transfer matrix placed higher attention on sequence variations of large regions, the effect of evolution is also manifested in recombination at the residue level or in small regions. To cope with this issue, we noticed that it is well studied the potential of protein language models to learn the intrinsic information from protein sequences, including evolutionary, functional, structural properties and those even remain unclear, thus building effective relationship networks based on sequence embeddings will contain details of various evolutionary relationships.

We developed a multi-step approach to construct and visualize the k-nearest neighbors (KNN) graphs from protein sequence embeddings (10). KNN algorithm calculates the distances between data points in the training set by metrics including Euclidean distance and Manhattan distance, based on the calculated distances, each point with K data points nearest are clustered. KNN graphs were constructed using the embedded protein sequences of interest and the outgroup sequence with the help of sklearn.neighbors package from the scikit-learn library, with a specified range of k-values and a distance threshold (17). The adaptive k-value, determined based on the threshold, ensures that only neighbors within a certain distance are considered, thus focusing on the most relevant connections, the choice of k-value depended on a balance between the clustering performance of the generated graph and the computational cost, we chose k = 15 in our KS/CLF analysis.

The resulting KNN graph was processed using the networkx library to extract all edges, representing connections between similar protein sequences based on their embeddings (18). These edges were saved for further analysis and visualization.

#### Evolution trajectory integration

KNN graph can display the clustering relationship of nodes while lacking information for the evolutionary trajectories. Therefore, we integrate the calculation of the module transfer matrix and edge weights/directions with the KNN graph. By extracting edges from the KNN graph and querying the weight matrix and edge direction results, we can obtain a directed graph with module transitions and assembly directions, providing insights into the evolutionary path simulations.

#### Nodes cluster and edge bundle analysis

We employed a hierarchical approach to cluster nodes and bundle edges in the KNN network (19)d. We performed hierarchical agglomerative clustering on the normalized PCA-reduced embeddings of the protein sequences while maintaining the distinction between KS and CLF categories. The FABF outgroup sequences were assigned to their nearest KS or CLF cluster based on sequence similarity. We determined the optimal clustering threshold as 30% of the maximum linkage distance, resulting in distinct clusters that each represent a group of closely related sequences within their respective KS or CLF categories. We then aggregated the edges between clusters, summing the weights of individual edges to create a condensed representation of the network. To enhance visual clarity, we implemented a custom edge bundling algorithm that processed bidirectional edges, retaining only the stronger connection when the weight difference exceeded 50% of the maximum weight between two clusters.

#### Root node prediction

To predict the root node of the integrated MAAPE network, we developed a scoring system based on its topological properties. For each node, we calculated a score as the ratio of its out-degree to its in-degree plus one, effectively measuring the node’s tendency to act as a source in the network. This approach is grounded in the assumption that ancestral sequences are more likely to have a higher proportion of outgoing edges, representing their role as evolutionary ancestors. The node with the highest score was designated as the predicted root.

#### Application of MAAPE for Gene Family Validation

To verify the reliability of the MAAPE method on other data sets, we selected two additional groups of bacterial proteins with previously studied evolutionary pathways as our research targets. One group consists of AlsR and AlsS, two divergently transcribed acetolactate synthase and acetoin biosynthesis transcriptional regulators, referred to as Group 1; The other group includes RadA, RadB, RecC, and Rad51, who play crucial roles in DNA repair and recombination-related functions across different domains of life, referred to as Group 2. We downloaded protein sequences for each gene from *Bacillus subtilis* in UniProt and performed BLAST searches against the non-redundant (nr) database using default parameters, top hit sequences from different domains of like for each search, resulting in 647 sequences for Group 1, 333 sequences for Group 2. We then conducted MAAPE analysis on these sequences and subsequently, performed node clustering and corresponding edge bundling, and found that the obtained evolutionary pathways were similar to the traditional consensus.

AlsS catalyzes the synthesis of compounds including acetoin, while AlsR, as a transcriptional regulator, controls the expression of the alsSD operon which includes AlsS (20). The relationship between AlsR and AlsS might be similar to that between KS and CLF, where the synthetic function existed first, and diverged to its regulatory partner in the purpose of diversity. Although in Figure 2b an archaeal AlsS is predicted as the root node, the AlsS_archaeon_3 group also has strong arrows indicating gene transfer towards the archaeal and bacterial groups of AlsR, which is consistent with the prediction.

**Figure 2.**
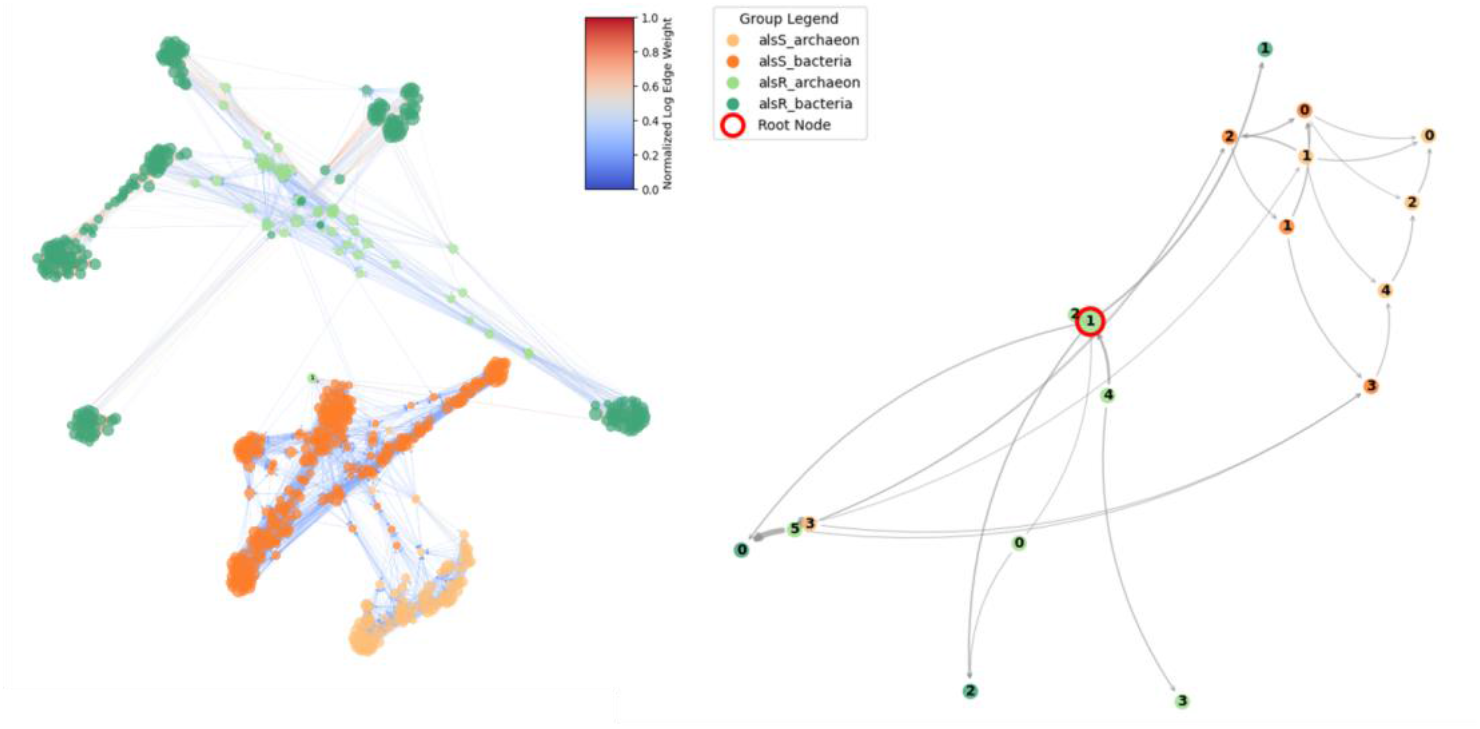
Visualization of evolutionary relationships between selected AlsR and AlsS sequences from archaeon and bacteria species. Using MAAPE network and its condensed representation. (a) The MAAPE network showing evolutionary relationships between sequences and their clustering information, where alsS_archaeon and alsS_bacteria (orange nodes) represent two archaeal and bacterial groups carrying the alsS gene, while alsR_archaeon and alsR_bacteria (green nodes) represent groups carrying the alsR gene. Directed edges indicate evolutionary trajectories, with edge colors representing connection strength according to the color bar. (b) A simplified backbone network derived from clustering and edge bundling, where nodes from the same group are consolidated and edge weights are bundled accordingly. This condensed visualization more clearly demonstrates the evolutionary paths between different groups, with the root node (marked in red circle) indicating the ancestral point of divergence.

The edges with greater weight are concentrated within the AlsR cluster rather than in the areas connecting clusters or within the AlsS cluster (21). This suggests the same ancestry and that horizontal gene transfer (HGT) occurs more frequently within the AlsR population. AlsS, being a metabolic enzyme, typically exhibits higher conservation across diverse species. In contrast, AlsR, functioning as a regulatory factor, demonstrates greater sequence plasticity. Clusters of AlsR are more dispersed than AlsS in Figure 2a, which also aligns well with this common view.

RadB, derived from *Methanococcus voltae*, is marked in orange as an outgroup. Due to the critical importance of genome repair functions, Genes in Group 2 are mainly conserved throughout species. In Figure 3a, we observe that the dispersion within each cluster is relatively small, and a clear evolutionary pathway is displayed. These four genes are believed to have originated from a common ancestry. RadB is considered to be the most ancient form in this gene family. Starting from RadB, this gene family differentiated for the adaptation to different domains of life: RadA primarily developed and was retained in archaea, RecA is widely present in bacteria, and Rad51 evolved in eukaryotes. In Figure 3b, RadB, along with a group of RecA and RadA, are positioned at the root of the network. This suggests that RadA and RecA might be intermediate forms that evolved from RadB, while Rad51 further differentiated from the other two genes after the emergence of eukaryotes. Although the main evolutionary path is vertical transmission, there may have been some HGT events between archaea and bacteria, which could explain the high-weight edge relationships between RadA and RecA near the root node in the figure.

**Figure 3.**
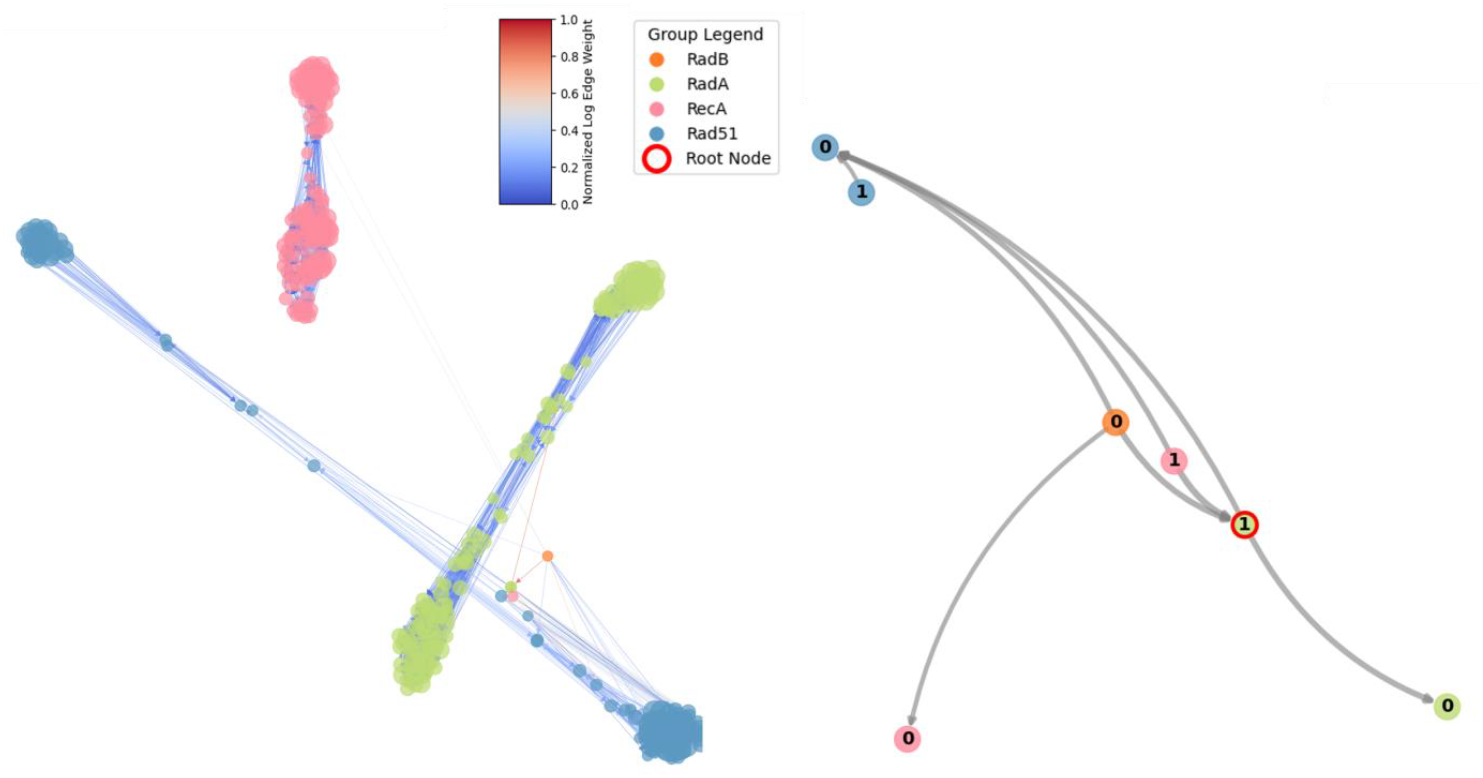
MAAPE network analysis of DNA repair protein families. (a) Full MAAPE network visualization showing evolutionary relationships between different DNA repair proteins. RadA and RadB (lime and orange nodes) represent two distinct rad gene families, while RecA and Rad51 (pink and blue nodes) represent additional DNA repair protein families. Edge colors indicate connection strength based on the color scale shown. (b) Condensed network representation after node clustering and edge bundling, with the root node (red circle) marking the ancestral point. This simplified view clearly illustrates the divergent evolutionary pathways between these DNA repair protein families, highlighting their relationships and potential functional diversification from a common ancestor.

Analysis of these two groups reveals that the shape and dispersion of the MAAPE plot are also crucial in providing clues about the evolutionary relationships. Symmetrical shapes in the plot suggest potential co-evolutionary relationships, a feature not observed in the other two analytical groups. Furthermore, tightly clustered nodes indicate more conserved sequence characteristics. The MAAPE plot’s topology offers additional insights into evolutionary dynamics, complementing traditional phylogenetic approaches. The symmetry observed here implies synchronized evolutionary processes, possibly due to functional interdependence or shared selective pressures. Conversely, the compact clustering of certain nodes underscores the high degree of sequence conservation, likely reflecting strong purifying selection on critical functional domains.

## Discussion

Conventional evolutionary analysis algorithms often employ alignment strategies that constructs guide trees based on pairwise sequence similarities (22). Phylogenetic algorithms inherently assume a fixed hierarchical evolutionary relationship among sequences, which simplifies the complex nature of evolutionary pathway (23). Such simplistic assumptions can lead to obscured alignments in regions where sequence identity falls below 30%, often referred to as the “twilight zone”, diminishing sensitivity and accuracy when dealing with proteins that exhibit significant sequence divergence yet maintain structural and functional conservation. In these low-similarity regions, the progressive alignment method may fail to account for intricate evolutionary events like horizontal gene transfer, convergent evolution, and compensatory mutations, resulting in less correct homologous pairings and misaligned residues. Moreover, the reliance on heuristic search strategies in many algorithms leads to compromises between computational efficiency and alignment precision, limiting their applicability to large and complex genomic datasets (24).

The persistent challenges of conventional phylogenetic methodologies call for new algorithms capable of dealing with enormous dataset while capturing accurate and nonlinear evolution network without compromising to misalignment. The ability of PLMs leveraging deep contextual and latent evolutionary patterns appears to perfectly tackle this problem (5). We have developed an evolutionary analysis method that dissects pLM embeddings across multiple window sizes to extract hierarchical subvectors. By assessing the similarities between these hierarchical subvectors, we construct a similarity matrix that not only represents the relationships between sequences but also indicates the direction of evolution from smaller to larger subvectors. This approach leverages the rich contextual representations generated by PLMs through sliding window approaches with stride of 1, capturing subtle evolutionary signals.

In all the datasets we benchmarked, the information entropy of vector segments derived from varying window sizes increases with segment length. However, beyond a length of 100, the rate of entropy increase significantly diminishes. This observation suggests that the 2560-dimensional embeddings generated by ESM-2 possess considerable compression capacity and demonstrate an efficient information preservation capability. Such properties indicate that ESM-2 embeddings effectively capture and condense the essential features of protein sequences, maintaining high informational integrity despite dimensional reduction. By embedding long sequences into uniformly sized vectors, ESM-2 facilitates the handling of extensive and diverse protein data without the computational burden typically associated with variable-length sequences. Furthermore, the uniform vector length enables the application of advanced dimensionality reduction techniques, such as Principal Component Analysis (PCA) or t-distributed Stochastic Neighbor Embedding (t-SNE), to further decrease computational load and enhance the efficiency of downstream analyses (13).

We further validated our evolutionary analysis method on datasets with established evolutionary relationships, including AlsS/R protein and a group of DNA repair protein families, achieving good performance. Instead of traditional hierarchical phylogenetic trees, our approach generates spatial relationship network graphs that elucidate the intricate relationships between different protein sequences. These network graphs encapsulate a multitude of informational dimensions, providing a comprehensive view that extends beyond the phylogenetic trees. In these validations, the predominant pathways align consistently with known evolutionary relationships, demonstrating the method’s accuracy in capturing established phylogenetic trajectories. However, our approach also uncovers correlations that are not readily apparent in traditional hierarchical evolutionary trees. These additional associations highlight the capability of PLMs to detect subtle and complex evolutionary relationships that standard tree-based methods may overlook.

In evolutionary trees, branch lengths represent evolutionary distances based on the number of amino acid residue differences, offering a linear and abstract measure of evolutionary divergence. In contrast, our method utilizes Euclidean distances between sequence positions within a high-dimensional embedding space. This geometric representation not only offers a more intuitive visualization but also captures richer information regarding evolutionary distances. By mapping sequences into a spatial framework, we can reveal complex evolutionary patterns and relationships that are often obscured in tree-based algorithms.

Overall, our PLM-based evolutionary analysis method leverages the powerful embedding capabilities of protein language models to overcome the limitations of traditional phylogenetic approaches. By providing a spatial and information-rich framework for visualizing evolutionary relationships, our method enhances both the accuracy and interpretability of evolutionary studies, offering significant advancements for fields such as protein engineering, functional genomics, and the comprehensive understanding of evolutionary mechanisms.

While we have observed that embeddings can be dissected into segments that retain specific informational properties, the precise functionalities and the extent of information conveyed by each sub-vector remain to be elucidated. Furthermore, the utilization of embeddings to address position-related challenges within the protein domain holds substantial developmental potential, our approach opens new avenues for solving complex spatial relationships buried in protein structures. This promising aspect requires extensive research to fully understand the capabilities of protein language models, ultimately advancing our ability to decode and manipulate the structural and functional property of proteins.

## Data availability

The authors declare that the data, materials and code supporting the findings reported in this study are available from the authors upon reasonable request.

## Author contributions

Zhiwei Qin and Heqian Zhang designed and supervised the research. Xiaoyu Wang and Jiaquan Huang performed the bioinformatic and established the algorithm. All authors analyzed and discussed the data. Xiaoyu Wang and Zhiwei Qin wrote the manuscript and all authors edited.

## Acknowledgements

This work was supported by the National Natural Science Foundation of China (32170079 to Z.Q., 32200035 to H.Z., and 32400235 to J.H.), the Natural Science Foundation of Guangdong (2024A1515012593 to Z.Q. and 2023A1515110175 to J.H.), Guangdong Talent Scheme (2021QN020100 to Z.Q.). The authors would like to thank the Interdisciplinary Intelligence Super Computer Center, Beijing Normal University, for High Performance Computing for access to computational resources.

